# Extensive neutralization against SARS-CoV-2 variants elicited by Omicron-specific subunit vaccine booster

**DOI:** 10.1101/2022.03.07.483373

**Authors:** Pai Peng, Chengqian Feng, Jie Hu, Chang-long He, Hai-jun Deng, Qinghong Fan, Jin Xiang, Guofang Tang, Mengling Jiang, Fengyu Hu, Feng Li, Kai Wang, Ni Tang, Xiaoping Tang, Ai-long Huang

## Abstract

The currently dominant variant of SARS-CoV-2 Omicron, carrying a great number of mutations, has been verified its strong capacity of immune escape in COVID-19 convalescents and vaccinated individuals. An increased risk of SARS-CoV-2 reinfection or breakthrough infection should be concerned. Here we reported higher humoral immune response elicited by Delta and Omicron variants after breaking through previous infection and cross-neutralization against VOCs, compared to the ancestral wild-type (WT) virus infection. To overcome the immune escape of Omicron, Omicron-specific vaccine was considered as a novel and potential strategy. Mouse models were used to verify whether Omicron-specific RBD subunit boost immune response by immunizing Omicron-RBD recombinant proteins. Three doses of Omicron-RBD immunization elicit comparable neutralizing antibody (NAb) titers with three doses of WT-RBD immunization, but the neutralizing activity was not cross-active. By contrast, two doses of WT-RBD with an Omicron-RBD booster increased the NAb geometric mean titers against Omicron by 9 folds. Moreover, an additional boost vaccination with Omicron-RBD protein could increase humoral immune response against both WT and current VOCs. These results suggest that the Omicron-specific subunit booster shows its advantages in the immune protection from both WT and current VOCs, and that SARS-CoV-2 vaccines administration using two or more virus lineages as antigens might improve the NAb response.

## Introduction

Since the coronavirus disease 2019 (COVID-19) pandemic caused by severe acute respiratory syndrome coronavirus 2 (SARS-CoV-2) began in 2019, it had experienced several waves driven by variants of this virus. At present, five variants including Alpha, Beta, Gamma, Delta and Omicron were designated as “variants of concern” (VOCs). As a dominate strain, even though the pandemic of Delta has lasted over one year, it has been replaced swiftly by Omicron (B.1.1.529) causing a new round of pandemic within a short time due to its rapid spread with a great deal of mutations^1^. More than 30 mutations have been accumulated in the spike (S) protein of Omicron variant, especially 15 of those occurs on receptor-binding domain (RBD), which is not only the vital binding site to the host receptor angiotensin-converting enzyme 2 (ACE2) for the entry of SARS-CoV-2, but also the key target of neutralizing antibodies produced by immune response and therapeutic antibodies. Spike mutation has been well documented to be correlated to its infectivity alteration and immune evasion^2-6^. The neutralizing activity of Omicron of sera has been suggested an extensive reduction from convalescents or vaccinees who received various types of SARS-CoV-2 vaccines ^3,7,8^.

Reduced neutralization elicited by infection or vaccination shows that Omicron has an increased risk of SARS-CoV-2 reinfection or breakthrough infection. In 31220 Norwegian households, secondary attack rate caused by Omicron was 25.1% (95% CI, 24.4%-25.9%)^9^. A study based on the Qatar national database suggests that the effective protection of previous infection against reinfection with the omicron variant was approximately 60%, which is lower than alpha, beta and delta variants (at approximately 90%)^10^. The vaccine effectiveness against Omicron after two BNT162b2 doses was 65.5% (95% confidence interval [CI], 63.9 to 67.0) at 2 to 4 weeks, and dropping to 8.8% (95% CI, 7.0 to 10.5) at 25 or more weeks^11^. Currently, an Omicron sub-variant BA.2 shows faster spread and similar resistance to immunity with high rate of breakthrough infection^12-14^. After breaking through previous immune protection by vaccine based on the ancestral wild-type (WT) virus, how the immune response elicited by Omicron breakthrough infection is needs to be delineated. It will give hints for the development of protective vaccine and vaccination strategies.

Due to continuous appearance of SARS-CoV-2 variants, reduced efficiency of existing vaccines accelerates the need of new vaccine strategies. A booster following primary vaccination series showed its potential efficiency of promoting high neutralizing activity^15^ and reducing symptomatic SARS-CoV-2 infection^16^, but booster shots displayed the failure in the breakthrough infection of some SARS-CoV-2 VOCs^17^. Moreover, simply additional boosters might not improve immune protection. A fourth-dose booster using the same antigen could not generate higher antibody titers than the third-dose vaccination and shows the low prevention against mild or asymptomatic Omicron infections and breakthrough infection^18^. Boosting with heterologous vaccines as one of the candidates, it has been proved to be safe and efficient immune response^**19-21**^. At present, several vaccination programs with heterologous vaccines have been approved in some countries. In view of the correlation of the immune escape of Omicron variant with its great number of mutations, Omicron-specific vaccines has been proposed. Omicron-specific mRNA vaccine booster could induce neutralizing response to Omicron itself but fail to previous VOCs^**22**^. As a booster with Omicron-matched DNA vaccines, increased width of immune response has been observed^**23**^. However, in macaque models, vaccination with Omicron-specific boosters do not increase neutralizing antibody (NAb) titers against Omicron and remain the equivalent levels of B cell response^**24**^. Of note, both two types of vaccines were designed according to full-length spike proteins. In consideration of the key role of SARS-CoV-2 RBD as the target of neutralizing antibodies, it has an important significance to verify whether Omicron-specific RBD subunit boost immune response after previous WT-RBD doses. Here we report immunogenicity and cross-reactivity of Omicron-specific RBD subunit proteins in mouse models to highlight the need of next generation of SARS-CoV-2 vaccines with variants-specific antigens.

## Results

In this study, twenty persons who were infected with Omicron or Delta after vaccination in each cohort were recruited, while 13 individuals previously infected with WT and unvaccinated were matched as a control cohort according to ages, sex and the time of sample collection of other two groups(Supplementary Table1). Delta breakthrough infections occurred 2.5-5 months (median 4.1months) after the last vaccine doses, while Omicron breakthrough infections occurred 3.4-6.6 months (median 5.2 months) after the last vaccine doses. Sera samples were collected at 3-4 time points within 50 days post symptom onset. Their anti-WT-RBD IgG binding antibody and neutralizing antibodies were determined by Enzyme-linked immunosorbent assay (ELISA) and the pseudotype-based neutralizing assay. Within the acute phase of COVID-19 infection, anti-WT-RBD IgG levels of sera in all three cohorts gradually raised to peak, then the antibody trends remained steady (Fig. 1A and Extended Data Fig. 1). As excepted, WT infection without additional immune protection, lower IgG titers were observed at the early stages of infection than Omicron or Delta breakthrough infection with their complicated histories of vaccination in different individuals. At the late stages of acute infection, anti-WT-RBD IgG binding antibodies reached to comparable levels among three cohorts.

**Fig. 1.**
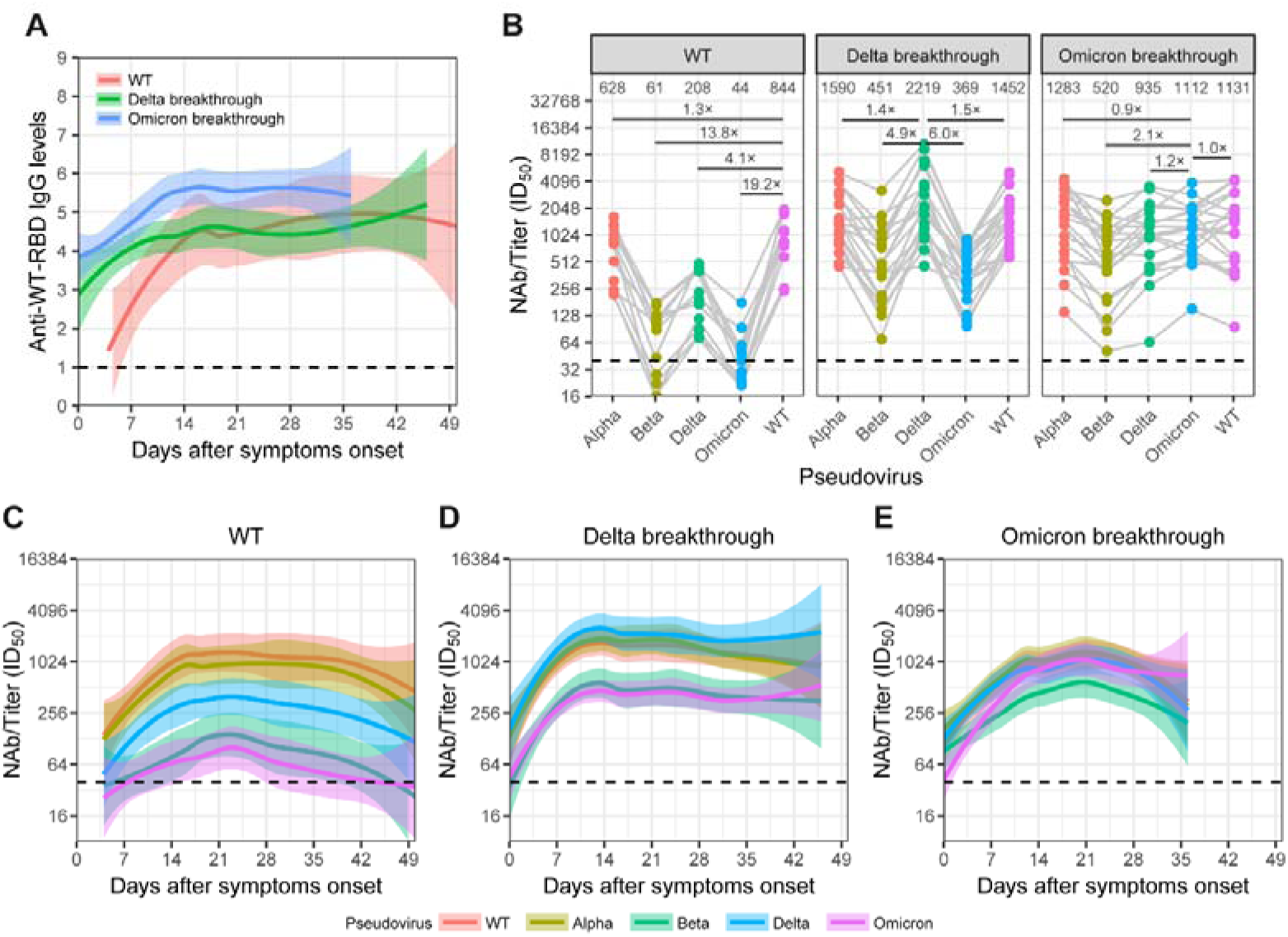
The characteristics of immune response elicited by the Omicron breakthrough infection. **A** dynamic change of anti-WT-RBD IgG binding antibodies in three cohorts: Omicron breakthrough infection or Delta breakthrough infection (n=20 for each cohort, sampled at 4 time points within 46 days after symptom onset), and WT infection without prior vaccination (sampled at 3 time points within 50 days after symptom onset). **B** Pseudovirus-based neutralizing assays were performed to test neutralizing activity of the sera against WT, Alpha, Beta, Delta and Omicron variants from three cohorts. **C-E** longitudinal observation of neutralizing antibody (NAb) titers of individuals experienced WT infection(C), Delta (D) and Omicron (E) breakthrough infection against WT, Alpha, Beta, Delta and Omicron pseudovirus. Statistical data analysis was performed using GraphPad Prism software. The half-maximal inhibitory dose (ID_50_) was calculated as NAb titers. Values above points indicate the geometric mean titers (GMTs). The threshold of ID_50_ detection was 1:40.

On the other hand, neutralization ability of breakthrough infection has been observed (Figure 1B, Extended Data Fig. 2). In the WT cohort, NAb titers of sera against four VOCs (Alpha, Beta, Delta and Omicron) are decreased by 1.3 folds, 13.8 folds, 4.1 folds and 19.2 folds, respectively, compared to the NAb titers against WT itself. In the Delta cohort, except for high NAb titers against Delta itself, NAb titers of sera against WT, Alpha, Beta and Omicron are decreased by 1.5 folds, 1.4 folds, 4.9 folds and 6.0 folds, respectively. In the Omicron cohort, except for high NAb titers against Omicron itself, NAb titers of sera against WT, Alpha, Beta and Delta are decreased by 1.0 folds, 0.9 folds, 2.1 folds and 1.2 folds, respectively, indicating the moderate cross-neutralization elicited by breakthrough infections.

**Fig. 2.**
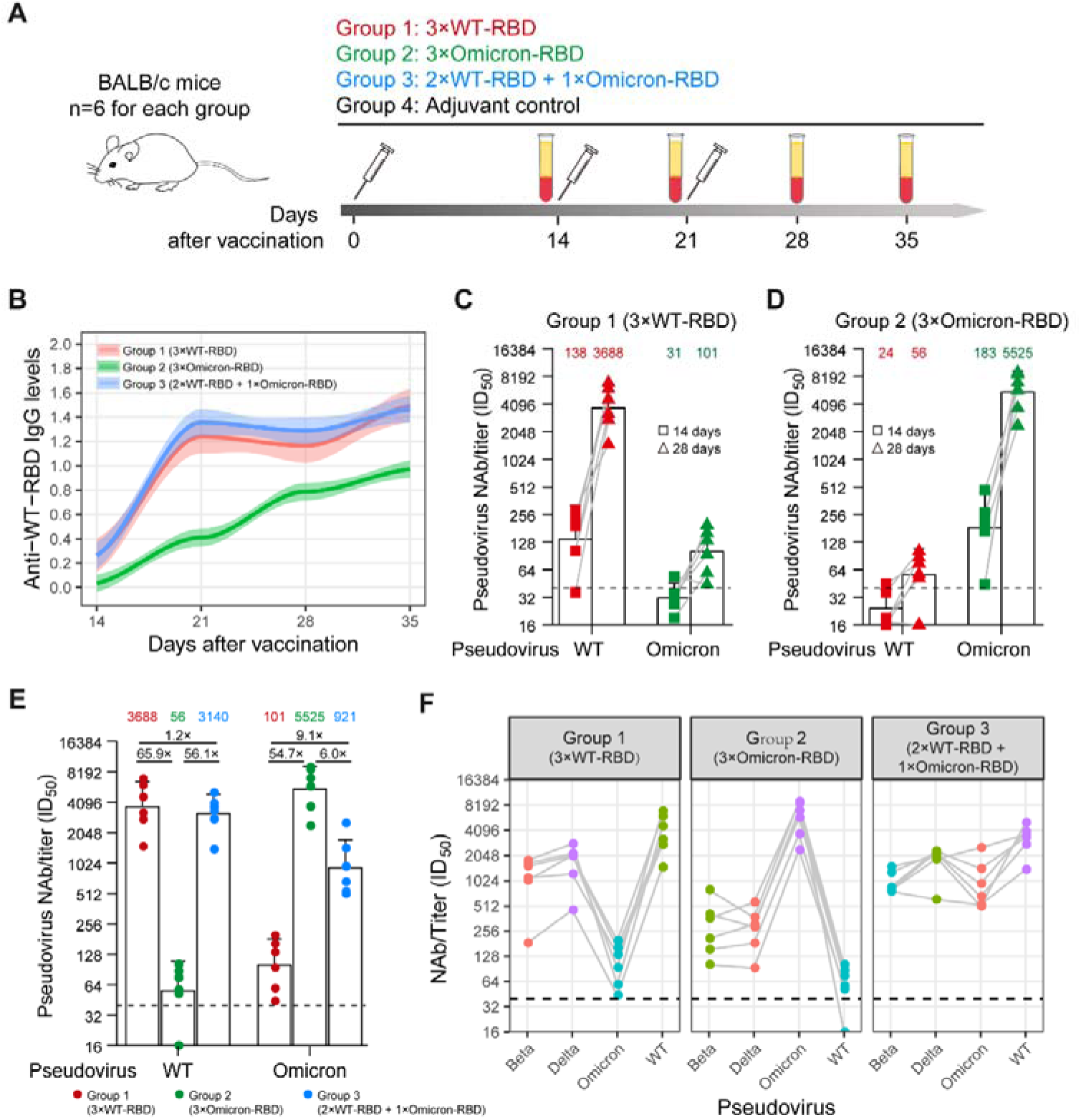
Humoral immune response elicited by Omicron-specific booster in mouse models. **A** Schematic of BALB/c mice immunization programs. 6 mice for each group were intramuscularly injected with the indicated recombinant proteins or the adjuvant as control. **B** dynamic change of anti-WT-RBD IgG binding antibodies in three groups within 35 days after the first dose of immunization. 1:16000 diluted sera were performed by ELISA. **C-D** Sera were collected and performed the pseudotype-based neutralizing assay to determine their neutralizing capacity to WT or Omicron pseudoviruses at 14 days (squares) and 28 days (triangels) after the first dose of immunization by WT-RBD (C) or Omicron-RBD (D) recombinant subunit proteins. **E** NAbs titers of sera against WT or Omicron from 3 groups collected at 28 days after the first dose of immunization. **F** Cross-neutralization of mice sera of 3 groups against WT, Beta, Delta and Omicron variants at 28 days after the first dose of immunization. Statistical data analysis was performed using GraphPad Prism version 8.0 software. Data on dot-bar plots is shown as GMT ± SEM with individual data points in plots. Values above points indicate the GMTs. The threshold of ID_50_ detection was 1:40.

During the follow-up visit, NAb titers among these three cohorts presents consistent dynamic changes with that of IgG antibodies (Figure 1C-E, Extended Data Fig. 3). WT cohort showed various degrees of neutralizing resistance to SARS-CoV-2 VOCs, especially to Beta and Omicron. In the Delta and Omicron cohorts, high neutralizing activity against themselves has been seen due to specific breakthrough infection. In the two breakthrough infection cohorts, even though NAb titers against WT induced by breakthrough infection did not exceed NAb levels in the WT cohort, sera displayed decreased neutralizing resistance against some variants. Taken together, compared with WT infection, breakthrough infection especially by Omicron could induce wide ranges and high levels of the humoral immune response to Omicron and other VOCs.

Widespread neutralizing activity of Omicron or Delta breakthrough infection against WT and other variants has been seen on the basis of the exposure of two antigens. Then, mouse models were used to evaluate whether previously ancestral vaccination or Omicron-specific immune response has the major contribution to the increased protection. BALB/c mice were distributed into 4 groups and immunized by two doses of SARS-CoV-2 RBD recombinant proteins subunit with 2-week interval plus one booster one week after the second dose (Figure 2A): Group1 immunized with 3 doses of WT-based RBD recombinant proteins (WT-RBD, Cat: K1516, Okaybio, China), Group2 immunized with 3 doses of Omicron-based RBD recombinant proteins

(Omicron-RBD,Cat: 40592-V08H121, Sinobiological, China), Group3 immunized with 2 doses of WT-RBD plus one dose of Omicron-RBD and Group4 immunized by the adjuvant (Cat: KX0210042, Biodragon, China) as the control.

Mice sera were collected and performed ELISA and the pseudotype-based neutralizing assay to determine their binding IgG levels and neutralizing effect on different VOCs. Among all these groups, two-week interval immunization induced the increased levels of anti-WT-RBD IgG binding antibodies (Fig. 2B). At Day 21 following the first dose of immunization, the third doses of RBD recombinant subunit proteins boost IgG antibody levels. Of note, higher anti-WT-RBD IgG levels have been shown in the sera elicited by three doses of WT-RBD proteins or two doses of WT-RBD proteins plus one dose of Omicron protein in Group1 and Group3, compared to Omicron-specific antibody response in Group2. It suggests partial cross-recognition of antibody response elicited by Omicron. On the other hand, booster by either WT-RBD or Omicron-RBD at the third dose of immunization produced the comparable levels of anti-WT-RBD IgG. The results are consistent with that of clinical samples we tested above.

Two-week interval with boosting by the third-dose immunization led to 26-fold increase (GMT from 138 to 3688) of NAbs against WT in Group1, while 30-fold increase (GMT from 183 to 5525) of NAb titers against Omicron variant in Group2 (Figure 2B-C). Omicron-specific NAb titers against Omicron in Group3 (GMT=5525) is comparable with WT-induced immune response (GMT=3688) to WT itself in Group1. It suggests that the immunogenicity of Omicron does not change. In contrast, sera from only WT-immunized mice in Group1 or only Omicron-immunized mice in Group2 do not neutralize other VOCs, except for itself (Figure 2D). However, it is noteworthy that Omicron-specific boosting after two-dose WT-RBD immunization can raise NAbs against Omicron by 9 folds. Subsequently, we tested cross-neutralization of these mice among three groups (Figure 2E). In Group1 only WT-RBD immunization showed high neutralizing capacity against WT itself and Delta, but weak neutralizing capacity against Omicron and Beta variants (3.7- and 2.4-fold decline, respectively). Likewise, in Group2, mice administrated with 3-dose of Omicron-RBD generated extremely high NAbs against Omicron variant itself but fail to neutralize other tested VOCs. Interestingly, in Group3, the Omicron-specific shot boost NAb titers not only against Omicron, but also against WT, Beta and Delta variants with GMT of 921, 3140, 962 and 1712, respectively. As NAb titers are found to be correlates with effective protection^25^, the data presented here suggest that Omicron-specific boosting will help to improve the immune protection from Omicron and that immunization by heterologous antigens will be beneficial to obtain wider protection against different SARS-CoV-2 variants.

## Discussion

Widespread immune escape in COCID-19 convalescents and vaccinees has been reported extensively. Furthermore, the fast transmission of Omicron and a surge of Omicron infected cases indicated the high risk of reinfection and breakthrough infection^17,26^. In a study about the influenza vaccine, authors suggest that prior infection enhances antibody responses to inactivated vaccine and is important to attain protective antibody titers^27^. In the Delta breakthrough infection after fully vaccination, 31-fold higher neutralizing antibody titers against the SARS-CoV-2 delta variant than vaccinees without infection was observed^28^. To evaluate the effect of prior vaccination on breakthrough infection, we analyzed the characteristics of humoral immune response elicited by Omicron variant after breakthrough infection. Compared to the previous infection with the ancestral strain WT, NAb titers against several variants from individuals infected with Delta or Omicron after breaking through the early immune protection generated by vaccines is significantly wider. Recent studies reported the consistent result that vaccination followed by breakthrough Omicron infection improved cross-neutralization of VOCs^29,30^, while neutralizing capacity of the unvaccinated individuals, which is triggered by Omicron, do not cross-neutralize other variants. These results suggest that prior immunity induced by vaccines will be beneficial to overcome the high neutralization resistance of Omicron^31^.

In light of extensive neutralization observed in Omicron breakthrough infection, we sought to understand the respective contribution of prior vaccination and Omicron-specific immunogenicity to this to establish more efficient immune protection. Therefore, we used immunized animal with WT-RBD and Omicron-RBD recombinant proteins to exhibit immune response elicited by Omicron-specific booster and heterologous antigens. Compared to NAb titers against WT pseudovirus produced by 3-dose of WT-RBD, Omicron-RBD alone can induce comparable NAbs against Omicron pseudovirus itself. It indicates that in mouse models, Omicron-RBD has the similar immunogenicity with WT-RBD. Furthermore, Omicron-RBD booster following two-dose of WT-RBD can induce 9-fold higher levels of NAbs against Omicron pseudovirus than the WT-RBD booster. We showed that Omicron-RBD boost following primary series could produce wider protection against the SARS-CoV-2 WT strain and circulating variants, which is consistent with what we observed above in Omicron breakthrough infection.

However, Omicron-mRNA boost in vaccinated macaques has not displayed significantly different NAb titers and B cells response^24^. This could be due to different immunization intervals and antigen epitopes. The time interval between vaccination and infection has been shown significant correlation with the potency of Omicron-neutralizing antibodies^32^. In our animal models, there is only 7-day interval between primary series (two doses) and booster. By contrast, immunization by Omicron-RBD subunit proteins shows the advantage that produce more NAbs to specially target against Omicron variant, while mRNA vaccine targeting full-length spike protein may produce more irrelevant antibodies, instead of targeting Omicron RBD^33^.

Our results indicate that heterologous antigens with various epitopes, which is different from single antigen as we have been vaccinated, may help to improve the height and width of NAb activity^34,35^. Except the booster vaccination strategies, A “bivalent” lipid nanoparticle (LNP) mRNA vaccine containing both Omicron and Delta RBD-LNP in half dose has been observed cross-neutralization against WT and three SARS-Cov-2 variants^36^. Multivalent vaccines could be alternative choice for the future development of SARS-CoV-2 vaccines and vaccination programs.

There are several limitations in our study. Due to limited participants with Omicron or Delta breakthrough infection were included in our study, the correlation of clinical characteristics with antibody response cannot be analyzed. Unvaccinated individuals who were infected with Omicron had not been recruited, but Omicron-specific immune response was observed in mouse models. Intramuscular injection in mouse models is not be completely equivalent to natural infection, hamster models could be used for virus challenge for further study.

Collectively, our data provides hints that the current booster vaccinations using WT-RBD protein or WT-S mRNA vaccine may be less efficient in preventing infections with the Omicron variant. Our results support the hypothesis that an additional boost vaccination with Omicron-RBD protein could increase humoral immune response against both WT and current VOCs.

## Acknowledgements

We acknowledge funding support from the China National Natural Science Foundation (grant no. U20A20392), the 111 Project (No. D20028), the Emergency Project from the Science & Technology Commission of Chongqing (cstc2020jscx-fyzx0053), Emergency Key Program of Guangzhou Laboratory (No. EKPG21-29 and EKPG21-31), Zhongnanshan Medical Foundation Of Guangdong Province No. ZNSA-2021004, China Postdoctoral Science Foundation (2021M693924), Natural Science Foundation of Chongqing, China (cstc2021jcyj-bshX0115), Chongqing Postdoctoral Science Special Foundation (2010010005216630) and the Research Fund Program of the Key Laboratory of Molecular Biology for Infectious Diseases, CQMU (No. 202105).

## Author Contributions

A.H., X.T., N.T., K.W., F.L, F.Y. and P.P. developed the conceptual ideas and designed the study. P.P., J.H., H.D. and C.H. performed the experiments and statistical analysis. C.F. provided the essential assistance through experiments. Q.F., G.T. and M.J. were responsible for sample collection. All authors provided scientific expertise and the interpretation of data for the work. P.P. drafted the manuscript. All authors contributed to critical revision of the manuscript for important intellectual content. All authors reviewed and approved the final version of the report.

## Materials and Methods

### Patients and samples

We enrolled 53 patients who had been identified to be previously infected with SARS-CoV-2 at the Eighth People’s Hospital of Guangzhou from January 2020 to January 2022. Thirteen patients infected with the SARS-COV-2 wild-type virus strain, 20 individuals infected with the SARS-COV-2 Delta virus after vaccination, and 20 patients infected with the SARS-COV-2 omicron virus after vaccination were included in our study. All infections were confirmed by q-PCR and sequenced to identify the genotype. The collection of all samples obtained the consent from subjects according to the protocols approved by the Ethics Review Board of the Eighth People’s Hospital of Guangzhou Institutional Review Board. Plasma was isolated from blood samples within 2h after collection according to the following steps: (1) Patient sera were heat incubated for inactivation at 56°C in water bath for 30 min; (2) centrifugation at 3000 rpm for 15 min, followed by transferring to new tubes; (3) Store at -80°C for further use.

### Mouse models and study design

8-week-old female BALB/c mice (6 mice per group) were provided by the Laboratory Animal Center of Chongqing Medical University (SCXK (YU) 2018-0003). Recombinant WT-RBD (Cat: K1516, Okaybio, China) and Omicron-RBD protein (Cat: 40592-V08H121, Sinobiological, China) as antigens for immunization were diluted with PBS, then mixed with an equal volume of QuickAntibody™-Mouse 3W adjuvant (Cat# KX0210042, Biodragon, China) and completely emulsified by a syringe. Each mouse was intramuscularly injected with 100 μl the antigen/adjuvant mixture. Serum samples were collected from tail tips before each vaccination and at 28 days after the first injection, then measure the antibody titers by ELISA and pseudovirus neutralization assay.

### Enzyme-linked immunosorbent assay (ELISA)

The recombinant RBD proteins derived from SARS-CoV-2 Wild-type (WT-RBD, Cat: K1516, Okaybio, China) and Omicron strains (Omicron-RBD, Cat: 40592-V08H121, Sinobiologial) were coated on 96-well microtiter plate (100ng/well) at 4°C overnight. After blocking with 5% skim milk powder and 2% BSA in PBS for 2 hours at room temperature, sera of enrolled patients were diluted and added into the plates and incubated at 37°C for 1 hour. After washing, wells were incubated with goat anti-mouse (Cat: ab6789, abcam, UK)/human (Cat: ab97225, abcam, UK) IgG-Horseradish peroxidase (HRP) antibody (1:10000 dilution) for 1 hour at 37°C. TMB substrate was added and incubated for 15 minutes at 37°C for color development. Reactions were stopped with stop solution, and the absorbance was determined at 450 nm using a microplate reader (Biotek, USA).

### Pseudovirus neutralization assay

For the neutralization assay, 50μL pseudoviruses of SARS-CoV-2 (Alpha, Beta, Delta, Omincron, D614G), equivalent to 3.8×10^4^ vector genomes, were incubated with serial dilutions of sera samples (dilutions of 1:40, 160, 640, 2560) from patients or mice for 1h at 37°C, then added the mixture into the 96-well plates seeded with 293T-ACE2 cells(1.6 × 10^4^ cells/well). The cells were refreshed with DMEM medium 8 h post-infection. Cells were lysed by 30 μl lysis buffer (Promega, Madison, WI, USA) at 72 h post-infection to measure RLU with luciferase assay reagent (Promega, Madison, WI, USA) according to the product instruction. Neutralization inhibition rate was calculated using GraphPad Prism 8.0 software (GraphPad Software, San Diego, CA, USA). The titers of neutralizing antibodies were calculated as 50% inhibitory dose (ID50).

### Ethics statement

Animal studies were approved by and conducted in compliance with the Committee on the Ethics of Animal Experiments of the Institutional Animal Care and Use Committee at the Laboratory Animal Center of Chongqing Medical University.

