## Supplementary Table 1 for "Extensive neutralization against SARS-CoV-2 variants elicited by Omicron-specific subunit vaccine booster"

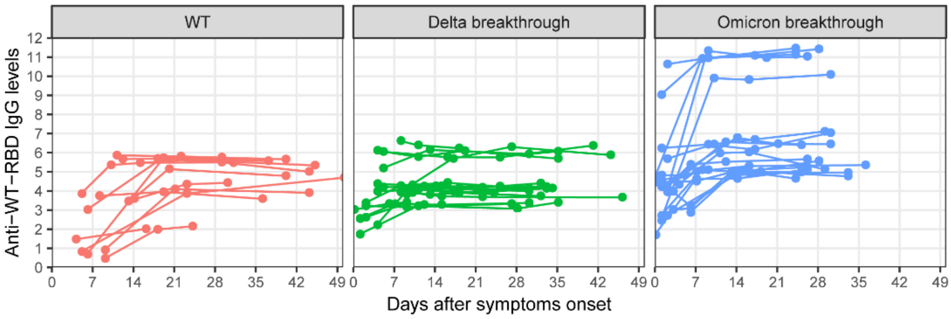


**Extended Data Fig. 1 Dynamic changes of anti-WT-RBD IgG binding antibodies in each individual of three cohorts.** A longitudinal observation of IgG antibody response to SARS-CoV-2 during the acute phase of wild-type (WT) infection, Delta and Omicron breakthrough infection.

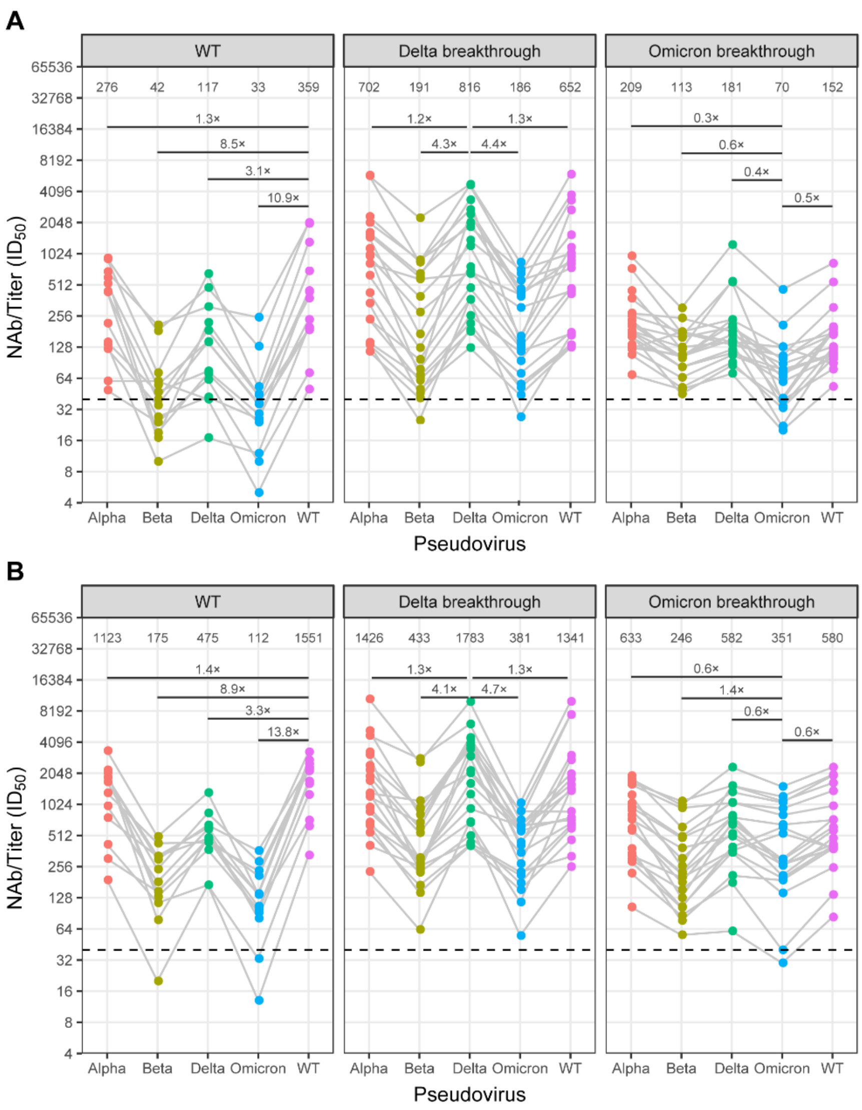


**Extended Data Fig. 2** Cross-neutralization against WT, Alpha, Beta, Delta and Omicron pseudoviruses in cohorts experienced WT infection, Delta or Omicorn breakthrough infection at the different time points.

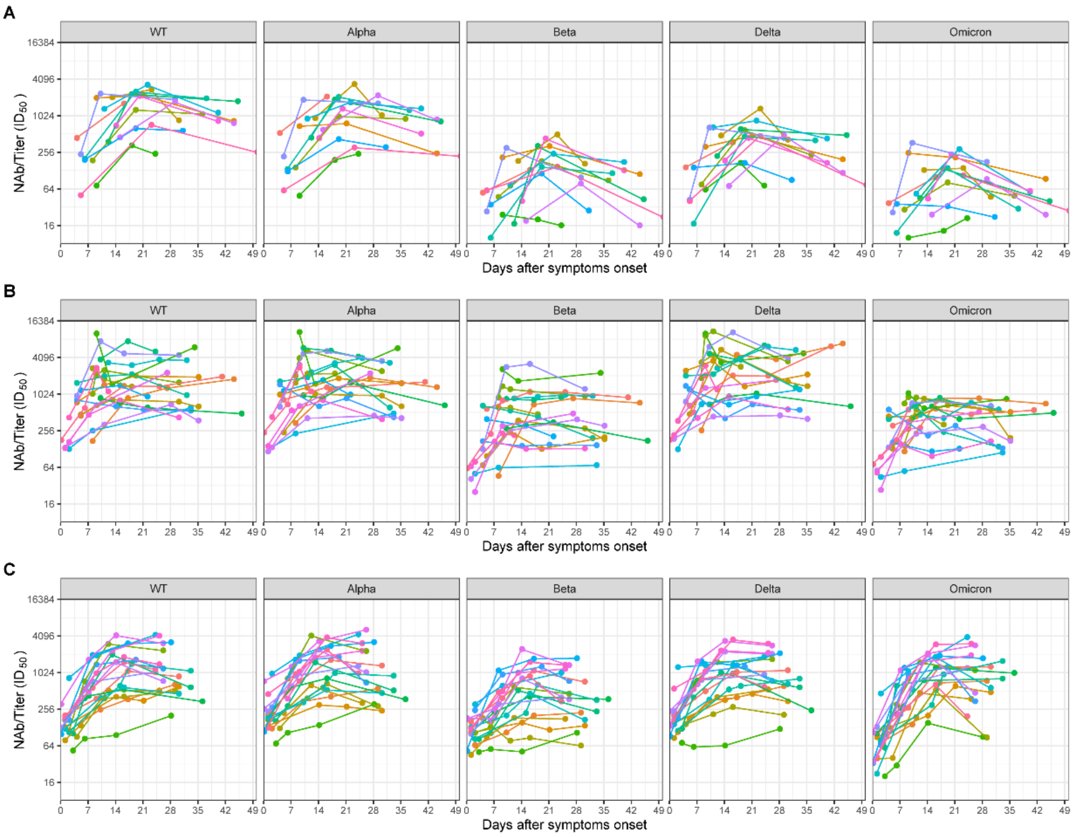


**Extended Data Fig. 3** Longitudinal observation of neutralizing antibody (NAb) titers of individuals experienced WT infection, Delta and Omicron breakthrough infection against WT, Alpha, Beta, Delta and Omicron pseudoviruses.

### Supplementary information

**Supplementary Table 1. Clinical characteristics of 53 individuals infected with SARS-CoV-2 wild-type strain, Delta or Omicron variants in this study.**

| Characteristics | COVID-19 cases | | | | P values | | |
|---|---|---|---|---|---|---|---|
| | Total (n=53) | Wildtype (n=13) | Delta (n=20) | Omicron (n=20) | W vs. D | W vs. O | D vs. O |
| **Age (mean±sd)** | 41.6±13.2 | 45.8±14.9 | 36.5±9.7 | 44.0±14.0 | 0.041 | 0.102 | 0.102 |
| **Sex** | | | | | | | |
| Male (n, %) | 34 (64.2%) | 5 (38.5%) | 13 (65.0%) | 16 (80.0%) | 0.169 | 0.027 | 0.480 |
| Female (n, %) | 19 (35.8%) | 8 (61.5%) | 7 (35.0%) | 4 (20.0%) | | | |
| **BMI** | 23.8 (21.0-25.5) | - | 22.6 (19.9-25.3) | 23.8 (21.7-25.5) | - | - | 0.425 |
| **Comorbidities (n, %)** | | | | | | | |
| Hypertension | 4 (7.5%) | 3 (23.1%) | 0 (0.0%) | 1 (5.0%) | 0.052 | 0.276 | 1.000 |
| Cardiovascular disease | 1 (1.9%) | 1 (7.7%) | 0 (0.0%) | 0 (0.0%) | 0.394 | 0.394 | 1.000 |
| Diabetes | 2 (3.8%) | 2 (15.4%) | 0 (0.0%) | 0 (0.0%) | 0.148 | 0.148 | 1.000 |
| Chronic liver disease | 2 (3.8%) | 1 (7.7%) | 1 (5.0%) | 0 (0.0%) | 1.000 | 0.394 | 1.000 |
| Chronic kidney disease | 1 (1.9%) | 1 (7.7%) | 0 (0.0%) | 0 (0.0%) | 0.394 | 0.394 | 1.000 |
| Thyroid disease | 1 (1.9%) | 0 (0.0%) | 0 (0.0%) | 1 (5.0%) | 1.000 | 1.000 | 1.000 |
| Any chronic disease | 7 (13.2%) | 5 (38.5%) | 1 (5.0%) | 1 (5.0%) | 0.025 | 0.025 | 1.000 |
| **Symptoms (n, %)** | | | | | | | |
| Fever | 33 (62.3%) | 9 (69.2%) | 14 (70.0%) | 10 (50.0%) | 1.000 | 0.310 | 0.333 |
| Cough | 29 (54.7%) | 4 (30.8%) | 17 (85.0%) | 8 (40.0%) | 0.003 | 0.719 | 0.008 |
| Expectoration | 24 (45.3%) | 3 (23.1%) | 14 (70.0%) | 7 (35.0%) | 0.013 | 0.701 | 0.056 |
| Pharyngalgia | 22 (41.5%) | 1 (7.7%) | 11 (55.0%) | 10 (50.0%) | 0.009 | 0.022 | 1.000 |
| Chest stuffiness | 5 (9.4%) | 1 (7.7%) | 4 (20.0%) | 0 (0.0%) | 0.625 | 0.394 | 0.106 |
| Nausea or Vomiting | 2 (3.8%) | 1 (7.7%) | 1 (5.0%) | 0 (0.0%) | 1.000 | 0.394 | 1.000 |

| | | | | | | | |
|---|---|---|---|---|---|---|---|
| Headache or Dizziness | 9 (17.0%) | 0 (0.0%) | 7 (35.0%) | 2 (10.0%) | 0.027 | 0.508 | 0.127 |
| Myalgia | 3 (5.7%) | 0 (0.0%) | 3 (15.0%) | 0 (0.0%) | 0.261 | 1.000 | 0.231 |
| Diarrhea | 2 (3.8%) | 1 (7.7%) | 1 (5.0%) | 0 (0.0%) | 1.000 | 0.394 | 1.000 |
| Fatigue | 6 (11.3%) | 0 (0.0%) | 4 (20.0%) | 2 (10.0%) | 0.136 | 0.508 | 0.661 |
| **Onset to admission, median days (median, IQR)** | 2 (1-3) | 4 (3-7) | 1 (0-3) | 1 (1-2) | 0.003 | 0.000 | 0.525 |
| **Onset to qPCR negative for SARS-CoV-2 (median, IQR)** | 16 (13-20) | 16 (12-20) | 14 (12-17) | 19 (16-23) | 0.349 | 0.168 | 0.003 |
| **Duration of hospitalization (median, IQR)** | 21 (17-26) | 21 (20-26) | 18 (15-32) | 21 (18-24) | 0.542 | 0.551 | 0.422 |
| **Laboratory results (median, IQR)** | | | | | | | |
| WBC | 5.06 (4.31-6.77) | 5.01 (4.06-5.64) | 5.30 (4.34-6.60) | 5.08 (4.52-7.40) | 0.548 | 0.418 | 0.565 |
| NEU | 3.18 (2.36-4.42) | 3.04 (2.00-3.50) | 3.21 (2.52-4.28) | 3.48 (2.41-5.88) | 0.316 | 0.207 | 0.301 |
| LYM | 1.13 (0.97-1.75) | 1.75 (1.13-2.07) | 1.12 (0.97-1.90) | 1.11 (0.83-1.30) | 0.246 | 0.008 | 0.317 |
| MONO | 0.48 (0.38-0.63) | 0.35 (0.29-0.42) | 0.45 (0.38-0.60) | 0.60 (0.52-0.71) | 0.028 | 0.000 | 0.050 |
| BASO | 0.02 (0.01-0.02) | 0.02 (0.01-0.02) | 0.02 (0.01-0.03) | 0.02 (0.01-0.02) | 0.152 | 0.883 | 0.183 |
| EOS | 0.04 (0.02-0.12) | 0.06 (0.04-0.10) | 0.08 (0.02-0.21) | 0.04 (0.03-0.06) | 0.684 | 0.273 | 0.369 |
| ALB | 44.7 (41.2-46.6) | 39.8 (38.3-41.6) | 45.0 (43.0-47.1) | 45.4 (43.5-46.7) | 0.000 | 0.000 | 0.745 |
| CRP | 9.00 (9.00-10.00) | 9.00 (9.00-10.00) | 9.00 (9.00-9.29) | 9.00 (9.00-9.00) | 0.511 | 0.360 | 0.868 |
| D-dimer | 0.21 (0.12-0.30) | 675.00 (605.00-747.50) | 0.18 (0.07-0.21) | 0.24 (0.15-0.30) | 0.002 | 0.000 | 0.034 |
| IL-6 | 5.73 (3.79-8.43) | - | 5.76 (4.86-8.75) | 5.70 (3.46-6.76) | - | - | 0.486 |
| PCT | 0.06 (0.03-0.08) | 0.06 (0.04-0.08) | 0.06 (0.05-0.08) | 0.02 (0.01-0.05) | 0.716 | 0.145 | 0.017 |
| LDH | 175.0 (153.5-203.0) | 180.0 (155.0-209.0) | 199.0 (173.0-211.0) | 165.5 (151.0-181.2) | 0.503 | 0.386 | 0.034 |
| T helper cell | 41.40 (30.48-43.72) | - | 35.60 (26.57-41.60) | 42.20 (35.55-45.22) | - | - | 0.083 |
| Suppressor T cell | 25.55 (20.65-30.82) | - | 27.75 (19.85-32.28) | 25.55 (21.97-29.00) | - | - | 0.767 |
| T helper cell/Suppressor T cell | 1.58 (1.11-1.96) | - | 1.28 (1.07-1.74) | 1.62 (1.55-1.96) | - | - | 0.184 |
| **Vaccination doses** | | | | | | | |
| Unvaccinated (n, %) | 13 (24.5%) | 13 (100.0%) | 0 (0.0%) | 0 (0.0%) | - | - | 0.231 |
| ≥3 dose (n, %) | 3 (5.7%) | - | 0 (0.0%) | 3 (15.0%) | | | |

| | | | | | | | |
|---|---|---|---|---|---|---|---|
| <3 dose (n, %) | 37 (69.8%) | - | 20 (100.0%) | 17 (85.0%) | | | |
| **Vaccination type** | | | | | | | |
| Unvaccinated (n, %) | 13 (24.5%) | 13 (100.0%) | 0 (0.0%) | 0 (0.0%) | | | |
| mRNA (n, %) | 6 (11.3%) | - | 3 (15.0%) | 3 (15.0%) | | | |
| Inactivated vaccine (n, %) | 29 (54.7%) | - | 14 (70.0%) | 15 (75.0%) | - | - | 1.000 |
| Co-inoculation | 1 (1.9%) | - | 0 (0.0%) | 1 (5.0%) | | | |
| Unknown (n, %) | 4 (7.5%) | - | 3 (15.0%) | 1 (5.0%) | | | |
| **Months of last vaccinated to the first sampling (median, IQR)** | 4.2 (2.6-6) | - | 4.1 (2.5-5) | 5.2 (3.4-6.6) | - | - | 0.082 |
| **Samples collected during follow-up (n, %)** | | | | | | | |
| ≤7 d.a.o | 44 (23.5%) | 5 (13.2%) | 15 (20.8%) | 24 (31.2%) | | | |
| 8-14 d.a.o | 46 (24.6%) | 8 (21.1%) | 22 (30.6%) | 16 (20.8%) | | | |
| 15-21 d.a.o | 32 (17.1%) | 9 (23.7%) | 12 (16.7%) | 11 (14.3%) | | | |
| 22-28 d.a.o | 22 (11.8%) | 4 (10.5%) | 8 (11.1%) | 10 (13.0%) | 0.131 | 0.009 | 0.362 |
| 29-35 d.a.o | 23 (12.3%) | 4 (10.5%) | 12 (16.7%) | 7 (9.1%) | | | |
| 36-42 d.a.o | 6 (3.2%) | 4 (10.5%) | 1 (1.4%) | 1 (1.3%) | | | |
| ≥43 d.a.o | 6 (3.2%) | 4 (10.5%) | 2 (2.8%) | 0 (0.0%) | | | |
